# Analysis of secondary metabolite gene clusters and chitin biosynthesis pathways of *Monascus purpureus* with high production of pigment and citrinin based on whole-genome sequencing

**DOI:** 10.1101/2022.01.31.478530

**Authors:** Song Zhang, Xiaofang Zeng, Qinlu Lin, Jun Liu

## Abstract

*Monascus* is a filamentous fungus that is commonly used for producing *Monascus* pigments in the food industry in Southeast Asia. While the development of bioinformatics has helped elucidate the molecular mechanism underlying secondary metabolite biosynthesis of *Monascus*, the biological information on the metabolic engineering of *Monascus* morphology remains unclear. In this study, we sequenced the whole genome of *Monascus purpureus* CSU-M183 strain by using combined single-molecule real-time DNA sequencing and next-generation sequencing platforms. The length of the genome assembly was 23.75 Mb in size with a GC content of 49.13% and 69 genomic contigs and encoded 7305 putative predicted genes. Furthermore, we identified secondary metabolite biosynthetic gene clusters and chitin synthesis pathways in the genome of the high pigment-producing *M. purpureus* CSU-M183 strain. And we confirmed that atmospheric room temperature plasma induced significant expression of the genes on *Monascus* pigments and citrinin biosynthetic gene cluster in *M. purpureus* CSU-M183 by RT-qPCR. These results provide a basis for understanding the secondary metabolite biosynthesis, the regulatory mechanisms of *Monascus* morphology, disrupting secondary metabolite biosynthesis in submerged fermentation, and the metabolic engineering of *Monascus* morphology.

## 1. Introduction

Fermented products of *Monascus* spp. have been widely used in the food and pharmaceutical industry for more than 2000 years.[1] As the secondary metabolite produced by *Monascus* spp., *Monascus* pigments (MPs) are a mixture of azaphilones mainly composed of three colors (yellow, orange, and red) pigments, which possess various bioactivities, such as antimicrobial, anticancer, anti-inflammatory, and anti-obesity.[2, 3] Nowadays, due to the potential risks of allergies, carcinogenesis, and teratogenesis of synthetic pigments, natural MPs are widely used as food colorants and are well recognized by consumers.[4, 5] In addition, MPs have other applications in the pharmaceutical, textile, and cosmetics industries. Traditionally, MPs are mainly produced by solid-state fermentation (SSF) with rice as the substrate for high pigment concentration.[6] Another method, submerged fermentation (SF) has high pigment production efficiency, an easy-to-control fermentation process, and avoidance of contamination, which makes SF attracted increasing attention and widely applied in the industrial production nowadays.[7, 8]

Different species of *Monascus* spp. have been isolated to study the biosynthesis of different secondary metabolites. In general, *M. fuliginosus*,[9, 10] *M. ruber[11, 12]* and *M. pilosus*[13–15] have a strong capacity of producing monacolin K. Additionally, due to its high efficiency to produce pigments, *M. purpureus* is one of the most predominant microorganisms for the efficient production of MPs.[16–18] With the development of bioinformatics of *Monascus*, the whole-genome sequencing (WGS) analysis has been used to reveal the chromosome evolution, regulatory mechanisms, and functional genes of *M. purpureus*, which lays the foundation for the production of secondary metabolites and biological researches. In 2015, Yang et al. published the first sequence information of *M*. *purpureus* YY-1, with a genome size of 24.1 Mb and a total of 7491 predicted genes. WGS analysis predicted the gene clusters related to pigment biosynthesis in *M. purpureus* YY-1 and explained the smaller size of the *M. purpureus* genome than that of related filamentous fungi, indicating a significant loss of genes.[19] Kumagai et al. reported the genome sequence information of the high pigment-producing *M. purpureus* GB-01 strain, with a genome size of 24.3 Mb and 121 chromosomal contigs.[20] Liu et al. identified the key genes (*ERG4*A and *ERG4*B) for ergosterol biosynthesis in *M. purpureus* LQ-6 (genome size: 26.8 Mb, 8596 protein-coding genes). Knocking out the *ERG4* gene improved the permeability of the cell membrane and secretion of intracellular pigments; it also changed the morphology of *M. purpureus* LQ-6 in SF broth.[21] Although many studies on the morphological changes of *Monascus* in SF have been performed,[22–24] the biological information on the metabolic engineering of *Monascus* morphology remains unknown.

Hyphae of filamentous fungi in SF mainly exist in three morphological forms, including free mycelia, mycelial pellets, and mycelial clumps,[25] and the biosynthesis of different target products tends to specific morphology. The mycelium pellet is the optimal morphology for glucoamylase production by *Aspergillus niger*, while the fermentation production of citric acid is more biased to the mycelial morphology.[26] The *ve*A gene globally regulates the propagation mode, mycelial growth, environmental tolerance, and secondary metabolites of fungi.[27, 28] Muller et al. disrupted the biosynthesis of chitin and changed the morphology of *Aspergillus oryzae* by regulating the transcription level of the chitin synthase gene *chs*B, and studied the relationship between morphology and α-amylase biosynthesis.[29] RNA interference technology has been applied to silence the expression of the chitin synthase gene *chs*4 in *Penicillium chrysogenum*, reducing the mutant growth rate, aggregation of dispersed hyphae into the mycelium, and increased penicillin production.[30] With the in-depth study of different phenotype mutants, it is found that the cell wall is an ideal target for morphological control. However, there are differences in the regulation of chitin synthase on morphology and the number of genes encoding chitin synthase in different fungi.[31] Additionally, the number of genes encoding chitin synthase and chitin biosynthesis pathway in *M*. *purpureus* is still unclear.

In this study, the whole genome of high-production pigments strain CSU-M183 was sequenced using the single-molecule real-time (SMRT, PacBioRS II) DNA sequencing and Illumina next-generation sequencing (NGS) platforms. The results showed a comprehensive prediction of biosynthetic gene clusters (BGCs) for secondary metabolites and the biosynthetic pathway of chitin in *M. purpureus* CSU-M183, providing new insights into morphological metabolism to regulate the biosynthesis of secondary metabolites.

## 2. Materials and methods

### 2.1 Fungal Strains, Culture Media, and Growth Conditions

*M. purpureus* CSU-M183 (CCTCC M 2018224, China Central for Type Culture Collection (CCTCC), Wuhan, China) was obtained using the atmospheric room temperature plasma (ARTP) mutation system from the parent strain *M. purpureus* LQ-6 (CCTCC M 2018600). Strains was cultivated on potato dextrose agar (PDA) and potato dextrose broth (PDB) medium at 30 °C in the dark for 7 days.

To prepare the inoculum, spores were transferred from PDA slants to submerged culture medium and washed with sterile distilled water, and then diluted to approximately 3 × 10^7^ spores/mL. The 10% (v/v) inoculum was transferred to the submerged culture medium and incubated 7 days. 10% (V/V) of the inoculum was transferred to 250ml shark flasks containing 45 ml liquid medium and incubated for 7 days in a rotary shaker with parameters set at 30 °C and 180 rpm, respectively.

### 2.2 DNA extraction

Mycelia were collected after centrifugation at 8228 ×*g* for 10 min and stored at −80 °C. Genomic DNA was extracted from mycelia using the *EasyPure*^®^ Genomic DNA Kit (TransGen Biotech, Beijing, China) according to the manufacturer’s protocol. The quantity, quality, and purity of the genomic DNA were measured using Nanodrop2000 systems and 0.8% DNase-free agarose gel electrophoresis.

### 2.3 WGS and assembling

The whole genome of the *M. purpureus* CSU-M183 strain was sequenced using SMRT sequencing technology of PacBioRS II, and the sequencing quality was improved using Illumina NGS platform. The sequencing library was constructed using the TruSeq™ Nano DNA LT Sample Prep Kit–Set A (Illumina, USA) and amplified using the TruSeq PE Cluster Kit (Illumina, USA).

For raw data polymerase reads after PacBioRS II sequencing, subreads were obtained by removing the low-quality or unknown reads, adapters and duplications. The filtered reads were assembled *de novo* using the Hierarchical Genome Assembly Process (HGAP) algorithm version 2.0.[32]

### 2.4 Gene prediction and annotation

AUGUSTUS[33] and SNAP[34] were performed to predict coding genes. Genome functional annotation was performed using BLASTP, as well as NCBI non-redundant (NR), SwissProt, and Protein Information Resource (PIR) protein databases. All predicted genes were classified according to the Kyoto Encyclopedia of Genes and Genomes (KEGG) metabolic pathways and Cluster of Orthologous Groups of proteins (COG).

### 2.5 Prediction of secondary metabolites

To predict secondary metabolite biosynthesis of strain *M. purpureus* CSU-M183, the BGCs of secondary metabolites were annotated using antiSMASH fungi version 5.1.0.[35]

### 2.6 RT-qPCR analysis

With β-Actin as the reference gene, the genes on the MPs and citrinin gene cluster were selected, and the expression of these genes was detected by qRT-PCR during the submerged fermentation of *M. purpureus* LQ-6 and *M. purpureus* CSU-M183, respectively. RT-qPCR was performed according to the method described by Liu et al.[21] The primers used in these analyses were listed in Table S1.

## 3. Results and discussion

### 3.1 Overview of WGS

In the previous study, we obtained a high pigment-producing *M*. *purpureus* CSU-M183 strain using the ARTP mutation system.[36] The morphological characteristics of *Monascus* are closely related to the production of secondary metabolites in SF. To further study the metabolic engineering of the morphology of *M*. *purpureus* CSU-M183, the WGS of strain *M*. *purpureus* CSU-M183 was carried out. The *M. purpureus* CSU-M183 genome sequence of 23.75 Mb was generated by assembling approximately 9.25 Gb raw data (353× coverage), which had a GC content of 49.13% and 69 genomic contigs (Table 1). The genome functional prediction and annotation identified 7305 protein-coding genes, with an average gene length of 1693 bp.

**Table 1.**
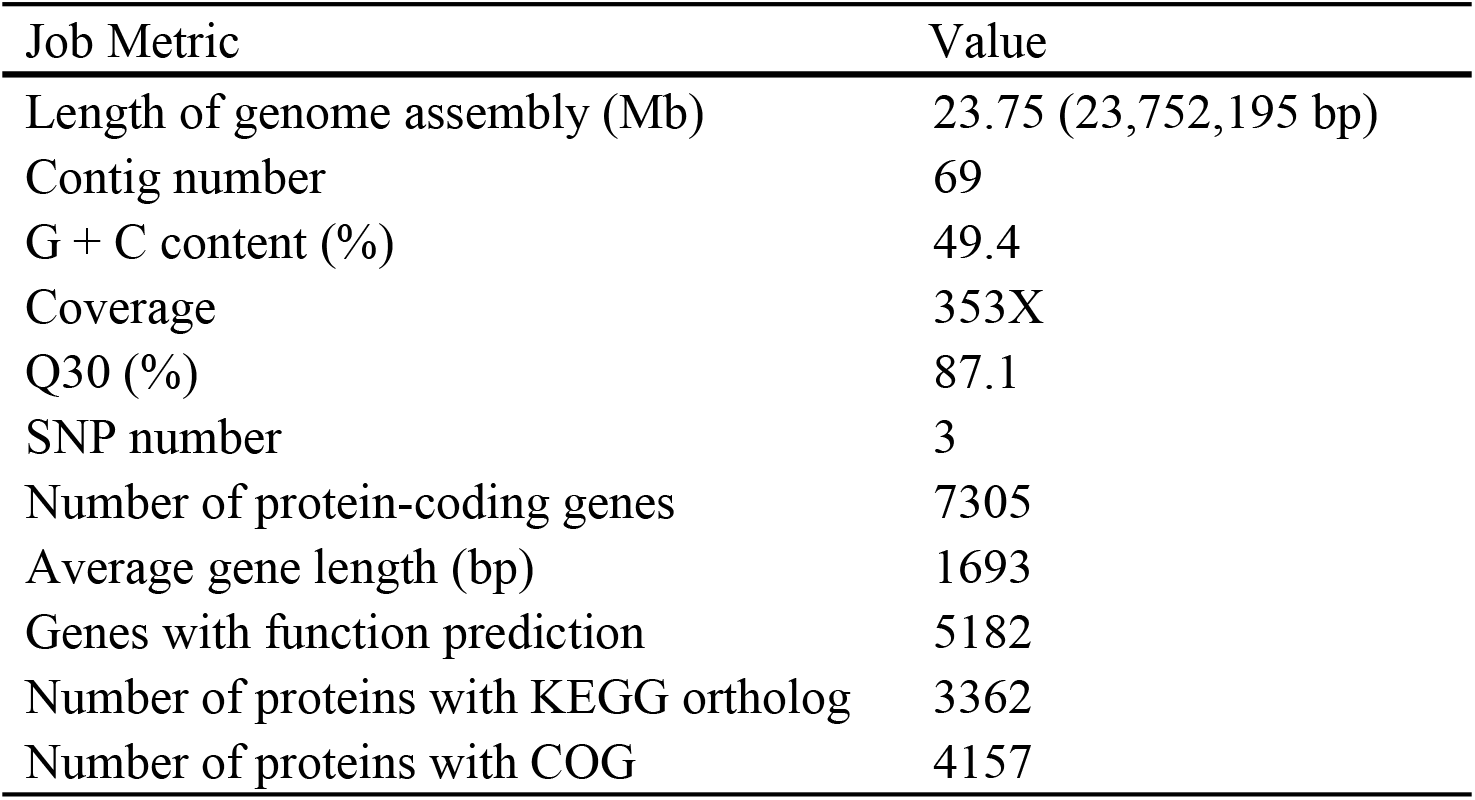
*M. purpureus* CSU-M183 genome general features

| Job Metric | Value |
| --- | --- |
| Length of genome assembly (Mb) | 23.75 (23,752,195 bp) |
| Contig number | 69 |
| G + C content (%) | 49.4 |
| Coverage | 353X |
| Q30 (%) | 87.1 |
| SNP number | 3 |
| Number of protein-coding genes | 7305 |
| Average gene length (bp) | 1693 |
| Genes with function prediction | 5182 |
| Number of proteins with KEGG ortholog | 3362 |
| Number of proteins with COG | 4157 |

To investigate the functions of the coding genes and metabolic pathways, all coding sequences (CDSs) were subjected to COG and KEGG analysis.[37] The COG database (http://www.ncbi.nlm.nih.gov/COG) classifies proteins by comparing all protein sequences in the genome.[38] In total, 4157 CDSs were allocated to COG categories (Fig. 1), with the maximum proportion of sequences related to “carbohydrate transport and metabolism” (8.52%), followed by “amino acid transport and metabolism” (7.82%), “translation, ribosomal structure and biogenesis” (6.90%), “posttranslational modification, protein turnover, and chaperones” (6.88%), and “energy production and conversion” (5.08%). Proteins that have not been fully identified in the genome of strain CSU-M183 were classified as “general function prediction only” (19.08%) and “function unknown” (4.81%) in COG categories.

**Fig. 1.**
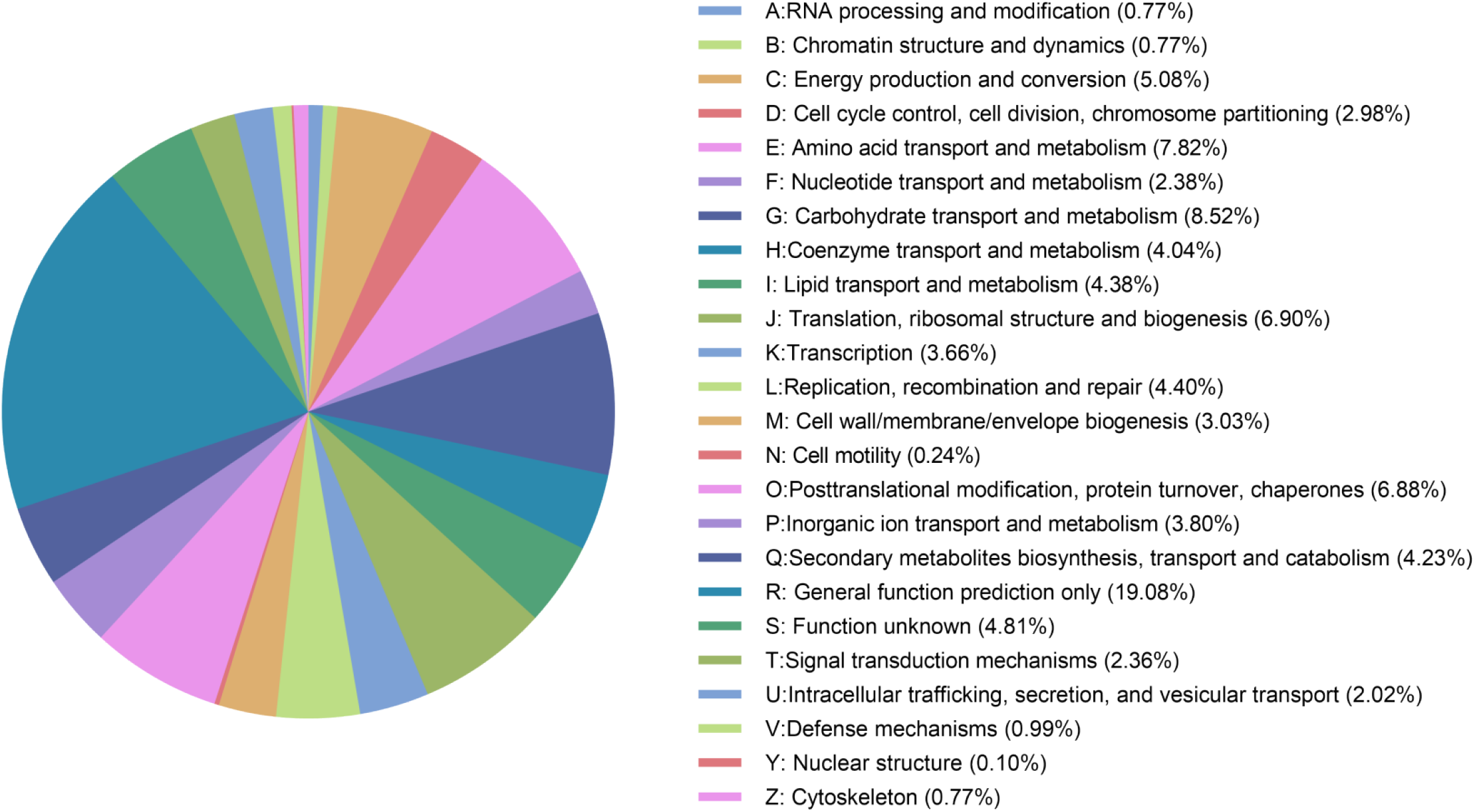
COG classification of predicted genes encoding proteins with annotated functions of *M. purpureus* CSU-M183 genome

KEGG enrichment analysis is essential for understanding the complex biological functions of genes in microorganisms, including metabolic pathways, genetic information transfer, and cytological processes.[39] Altogether, 3362 CDSs were allocated to five categories in the KEGG database, including “metabolism”, “cellular process” and “environmental information processing”, “genetic information processing”, and “organismal systems” (Fig. 2). Annotation results showed that “metabolism” is the main category of KEGG annotations (1375, 40.90%), followed by “genetic information processing” (707, 21.03%) and “organismal systems” (519, 15.44%). Moreover, CDSs were significantly enriched in “translation” (282), “carbohydrate metabolism” (279), “amino acid metabolism” (266) and “transport and catabolism” (259) subcategories, indicating that *M*. *purpureus* CSU-M183 had the strong ability of protein translation, carbohydrate utilization and energy conversion.

**Fig. 2.**
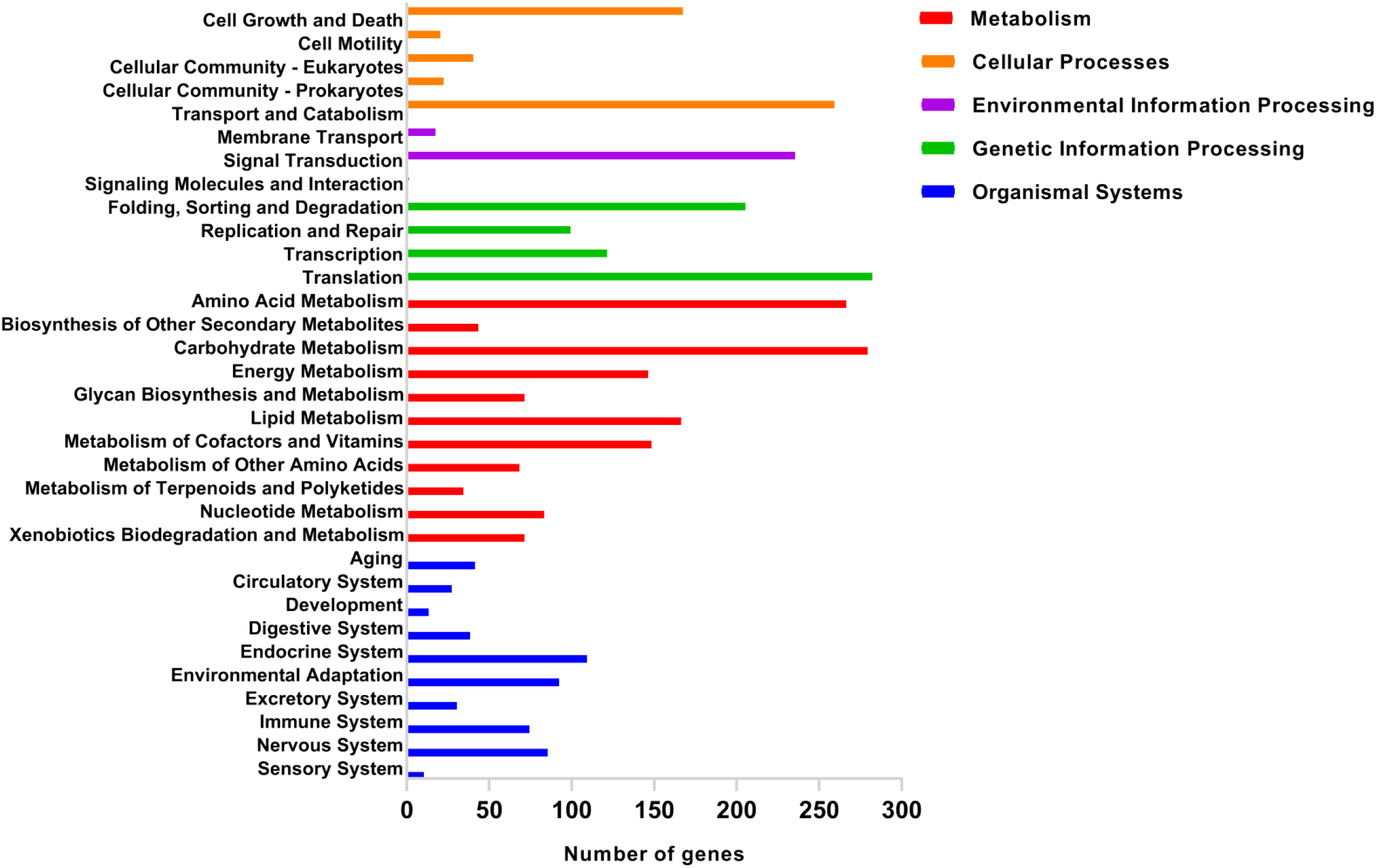
Enrichment analysis of KEGG pathways for predicted genes of the *M. purpureus* CSU-M183 genome. The y-axis represents the KEGG pathway and the x-axis denotes the number of genes

### 3.2 Identification of secondary metabolites BGCs

AntiSMASH is a widely used tool that can identify and annotate BGCs in bacterial and fungal genome sequences.[40] To further understand the biosynthesis of the secondary metabolites in strain *M*. *purpureus* CSU-M183, BGCs prediction of secondary metabolites were performed using antiSMASH fungi version 5.1.0. A total of 26 BGCs were detected, including terpene, non-ribosomal peptide synthetases (NRPS), type I polyketide synthases (T1PKS), β-lactone, and 18 putative gene clusters. After the database search with antiSMASH, a BGC (contig 000002F, gene g3398-g3411) for citrinin within the genome sequence of strain CSU-M183 was predicted (Fig. 3a), which was identical to the known citrinin BGC (GenBank accession number: AB243687.1, 21917 bp),[41] and the identity of homologous genes was 99%-100%, the predicted functions of the genes in citrinin BGC are listed in Table 2. Additionally, 81% of homologous genes were similar to those in citrinin BGC0001338, and 57% in citrinin BGC000894. A putative BGC responsible for the biosynthesis of MPs was identified in the genome of strain CSU-M183 with 41% of homologous genes showed similar to that in BGC0000027 (Fig. 3b), including 16 genes (contig 000001F, gene g1401-g1416) listed in Table 3. As shown in Table 3, the identity of homologous genes was considerably high, such as gene g1409 was 96.51% similar to *Mpig*A (Polyketide synthase), gene g1407 was 95.05% similar to *Mpig*C (Ketoreductase), and gene g1406 was 95.82% similar to *Mpig*D (Acyltransferase). Moreover, by using the known monacolin K BGC (GenBank accession number: DQ176595.1, 45000 bp) of *M*. *pilosus* as a reference,[13] no complete monacolin K BGC was detected in the genome sequence of *M. purpureus* CSU-M183 (Fig. 3c). All protein-coding genes in the genome sequence were analyzed by BLASTP, where gene g3061, g2167, g4491, g1403, g1402, g4228, g1429, g1395 were homologous to the genes mkA~mkI, respectively. However, these genes do not locat in a gene cluster in genome, and the homologous protein identities were low, especially gene g1403 (with *mokD* of *M*. *pilosus* with 25.00% identity) (Table 4). It has been reported that overexpression of *mokD* significantly enhanced the production of monacolin K by 200.8%, which illustrated this gene play a vital role in the synthesis of monacolin K.[24] These findings were similar to that of the parent strain *M. purpureus* LQ-6, which could not produce monacolin K.[42]

**Table 2.** Functional prediction of genes detected in the citrinin BGC of *M. purpureus* CSU-M183

| CDS | Length<br>(bp) | Product | KO |
| --- | --- | --- | --- |
| gene=g3398 | 1762 | Unnamed protein product |  |
| gene=g3399 | 1389 | Hypothetical protein, *orf7* |  |
| gene=g3400 | 2796 | *ctnD* | K00108 |
| gene=g3401 | 403 | Predicted protein |  |
| gene=g3402 | 3076 | Dehydrogenase, *orf1* | K13821 |
| gene=g3403 | 987 | citrinin biosynthesis oxygenase, *ctnA* |  |
| gene=g3404 | 940 | citrinin biosynthesis oxydoreductase,<br>*ctnB* |  |
| gene=g3405 | 7780 | citrinin polyketide synthase, *pksCT* |  |
| gene=g3406 | 1497 | citrinin biosynthesis transporter, *ctnC* |  |
| gene=g3407 | 829 | *ctnF* |  |
| gene=g3408 | 1498 | Hypothetical protein, *orf8* |  |
| gene=g3409 | 2116 | *ctnR* |  |
| gene=g3410 | 663 | *ctnG* | K01673 |
| gene=g3411 | 567 | *ctnG* |  |
| gene=g3412 | 3163 | *ctnI* |  |

**Table 3.** Functional prediction of genes detected in the MPs BGC of *M. purpureus* CSU-M183

| CDS | Length<br>(bp) | Product | Identity<br>(%) | KO |
| --- | --- | --- | --- | --- |
| gene=g1401 | 2388 | Fungal specific transcription factor, *MpigR* | 91.41 |  |
| gene=g1402 | 1108 | Alcohol dehydrogenase, *MpigE* | 95.39 |  |
| gene=g1403 | 819 | Amino oxidase/esterase, *MpigF* | 93.01 |  |
| gene=g1404 | 1395 | Amino oxidase/esterase, *MpigF* | 94.03 |  |
| gene=g1405 | 1027 | Dehydrogenase, *MpigH* | 97.66 |  |
| gene=g1406 | 1449 | Acyltransferase, *MpigD* | 95.82 | K22889 |
| gene=g1407 | 892 | Ketoreductase, *MpigC* | 95.05 | K11165 |
| gene=g1408 | 1719 | Citrinin biosynthesis transcriptional activator<br>CtnR | 95.48 |  |
| gene=g1409 | 8071 | Polyketide synthase, *MpigA* | 96.51 |  |
| gene=g1410 | 1032 | Peptidyl-prolyl cis-trans isomerase Cpr7 | 98.68 | K05864 |
| gene=g1411 | 573 | D-tyrosyl-tRNA(Tyr) deacylase, *MpigO* | 95.58 | K07560 |
| gene=g1412 | 1849 | TPA: CCR4-NOT transcription complex,<br>subunit 3 | 84.23 | K12580 |
| gene=g1413 | 634 | C<sub>2</sub>H<sub>2</sub> finger domain protein | 76.40 |  |
| gene=g1414 | 1531 | Dihydrolipoamide dehydrogenase | 84.17 | K00382 |
| gene=g1415 | 1665 | Fatty acid desaturase | 73.58 | K13076 |
| gene=g1416 | 1792 | Phosphoglucomutase | 68.45 | K01835 |

**Table 4.** Functional prediction of genes detected in the monacolin K BGC of *M. purpureus* CSU-M183 by NCBI-BLASTP.

| CDS | Length (aa) | Homologue | Homologue<br>Length (aa) | Identity |
| --- | --- | --- | --- | --- |
| gene=g3061 | 3945 | ABA02239.1_8<br>[gene=mkA] | 3075 | 37.19% |
| gene=g2167 | 2593 | ABA02240.1_1<br>[gene=mkB] | 2547 | 38.62% |
| gene=g4491 | 499 | ABA02241.1_9<br>[gene=mkC] | 524 | 80.53% |
| gene=g1403 | 273 | ABA02242.1_7<br>[gene=mkD] | 263 | 25.00% |
| gene=g1402 | 369 | ABA02243.1_6<br>[gene=mkE] | 360 | 39.17% |
| gene=g4228 | 1133 | ABA02245.1_4<br>[gene=mkG] | 1052 | 52.61% |
| gene=g1429 | 426 | ABA02246.1_3<br>[gene=mkH] | 455 | 78.10% |
| gene=g1395 | 570 | ABA02247.1_2<br>[gene=mkI] | 543 | 54.21% |

**Fig. 3.**
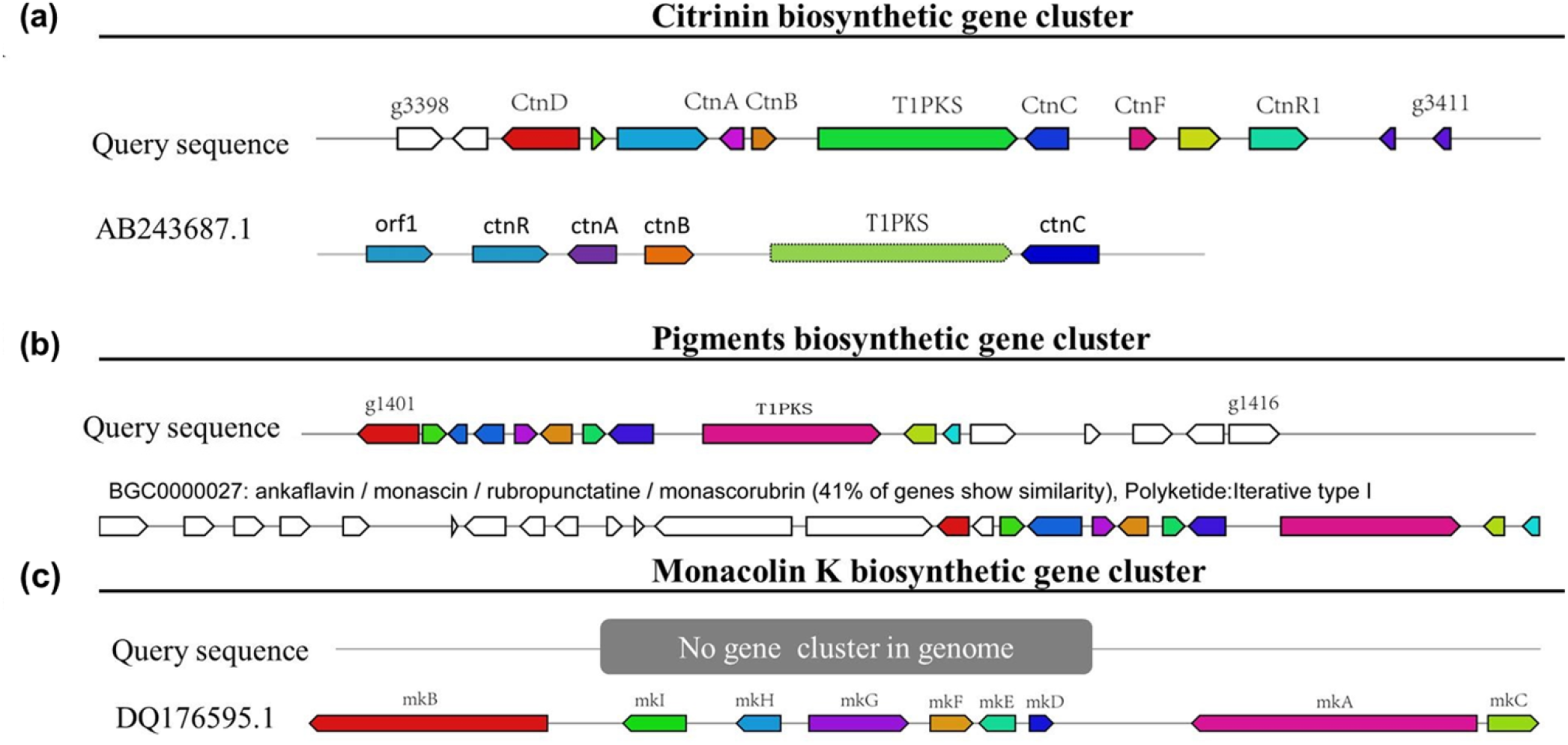
Schematic representation of predominant secondary metabolites BGCs in the genome sequence of *M. purpureus* CSU-M183. (a) CIT. (b) MPs. (c) Monacolin K

Due to various species of *Monascus* and large gaps in available biological information, the development of basic theoretical research on *Monascus* has been relatively slow. With the continuous development of sequencing technology and bioinformatics, breakthrough progress has been made in biosynthetic pathways of the secondary metabolites of *Monascus*,[19, 43, 44] among which MPs, citrinin and monacolin K are the most notable. However, studies on functional and comparative genomics—such as the annotation of unknown sequences, investigation of gene models, comparison of multiple sequence alignment analysis, and metabolic engineering of *Monascus* morphology, which can elaborate the relationship among secondary metabolite productivity, growth, and morphology in SF, are less.

The key gene PKS with 7838 bp responsible for citrinin biosynthesis was first identified from *M*. *purpureus* in 2005[45] and then five genes encoding Zn(II)_2_Cys_6_ transcriptional activator, membrane transporter, dehydrogenase, oxygenase, and oxidoreductase for citrinin biosynthesis were cloned.[46] Microbial PKSs have been mainly classified into three types –type I PKSs (modular type I PKSs and iterative type I PKSs), type II PKSs, and type III PKSs. The PKS responsible for citrinin biosynthesis belongs to the iterative type I PKSs, which contains putative domains for ketosynthase (KS), acyltransferase (AT), ketoreductase (KR), dehydratase (DH), enoyl reductase (ER), methyltransferase (MT), thioesterase (TE), and acyl carrier protein (ACP).[45] The KS domain catalyzes the condensation of precursors to extend the polyketone chain, whereas the AT domain selects the precursors, and the ACP domain makes covalent bonds between the precursors and intermediates, which are necessary for the functioning of most PKSs.[47] In 2012, the citrinin BGC with the length of 43 kb from *Monascus aurantiacus* was first published,[48] including 16 open reading frames (ORFs) for *ctn*D, *ctn*E, *orf*6, *orf*1, *ctn*A, *orf*3, *orf*4, *pks*CT, *orf*5, *ctn*F, *orf*7, *ctn*R, *orf*8, *ctn*G, *ctn*H, and *ctn*I, which are dramatically similar to those of the CIT BGC of strain *M. purpureus* CSU-M183. These results revealed high homology of citrinin BGC in *Monascus,* especially the key gene PKS. In 2012, a putative 53 kb MP BGC of *M*. *ruber* was first reported, which consisted of genes encoding PKSs, fatty acid synthases, regulatory factors, and dehydrogenase.[3] Xie et al. reported that gene *pigR* (gene g1401 in Table 3) is a positive regulatory gene in MPs biosynthesis pathway,[49] whereas gene *MpigE* (gene g1402 in Table 3) may be involved in the conversion of different MPs.[50] The genome size of *M. purpureus* was found to be smaller than that of related filamentous fungi, indicating a significant loss of genes.[19] A previous study reported that monacolin K cannot be produced due to the lack of monacolin K biosynthesis locus in some *M. purpureus* genomes.[51] After the prediction of monacolin K BGC in the genome of strain CSU-M183, we found that there was no complete monacolin K BGC in the strain CSU-M183, which was consistent with previous studies.[42, 51] Undoubtedly, the identification of BGCs has greatly facilitated the understanding of the biosynthetic pathways of secondary metabolites in *Monascus*, which can provide theoretical support for industrial production of *Monascus* secondary metabolites.

### 3.3 The genes’expression in MPs and citrinin gene cluster from ***M. purpureus*** CSU-M183 and ***M. purpureus*** LQ-6

After 7 days of SF, the MPs and citrinin yields of *M. purpureus* LQ-6 and *M. purpureus* CSU-M183 were 43.97 U/mL, 1.27 mg/L and 83.77 U/mL, 5.34 mg/L, respectively(Fig. 4a). To verify the effect of mutagenesis on the metabolism of MPs and citrinin, the relative expression levels of several key genes, *MpigA*, *MpigR*, *MpigC*, *MpigD*, *MpigE*, *MpigF*, *MpigG*, *MpigH*, *MpigI*, *MpigJ*, *MpigK*, *MpigL*, *MpigM*, *MpigP*, *MpigQ*, *cit S*, *cit A*, *cit B*, *cit C*, *cit D* and *cit E* were vestigated using RT-qPCR. As shown in Fig. 4b and 4c, the relative expression levels of *MpigM*, *MpigP*, *cit B*, *cit C*, *cit D* and *cit E* in *M. purpureus* CSU-M183 were extremely significant compared to that of *M.purpureus* LQ-6 at 4th day; while the relative expression levels of *MpigA*, *MpigJ*, *MpigK*, *cit S*, *cit C* in *M. purpureus* CSU-M183 were obviously higher than that of *M. purpureus* LQ-6 at 7th day. During the fermentation process, in addition to the extremely significant genes with relatively significant expression levels, the expression levels of other genes in the MPs and citrinin synthetic gene clusters in M183 were also very significant, such as *MpigC*, *MpigD* and *cit S* at 4th day. The production of MPs and citrinin is directly or indirectly related to the function of genes in their biosynthetic gene clusters, and the relative expression of genes can directly reflect the contribution of genes in the fermentation process. A number of studies have performed functional analysis of MPs and citrinin gene clusters, such as: inactivating *MpigA* in *M. ruber*, *Monascus* lost its pigment production ability, which proved that PKS was involved in pigment synthesis.[52] MrpigJ(encoded by *MrpigJ*, a homolog of *MpigJ*) and MrpigK(encoded by *MrpigK*, a homolog of *MpigK*) form two subunits of the specialized fungal FAS, which produce the fatty acyl portion of the side chain of MPs.[53] Moreover, MrpigM, as an o-acetyltransferase, synthesized an O-11 acetyl intermediate in Chen et al’s *Monascus* model, and knocking out *MrpigM*(the homolog of *MpigM*) blocked the pathway of pigment synthesis intermediate[53]. The inactivation of genes in citrinin biosynthesis gene cluster led to a significant decline on citrinin production, even lower than the detection level, such as knocking out *cit A*, *pksCT* and *cit B*.[41, 54] It indicates that the increase of MPS and citrinin production may be caused by the increase of gene expression level in gene cluster caused by ARTP mutation, and these genes are very important for the synthesis of MPS and citrinin.

**Fig. 4.**
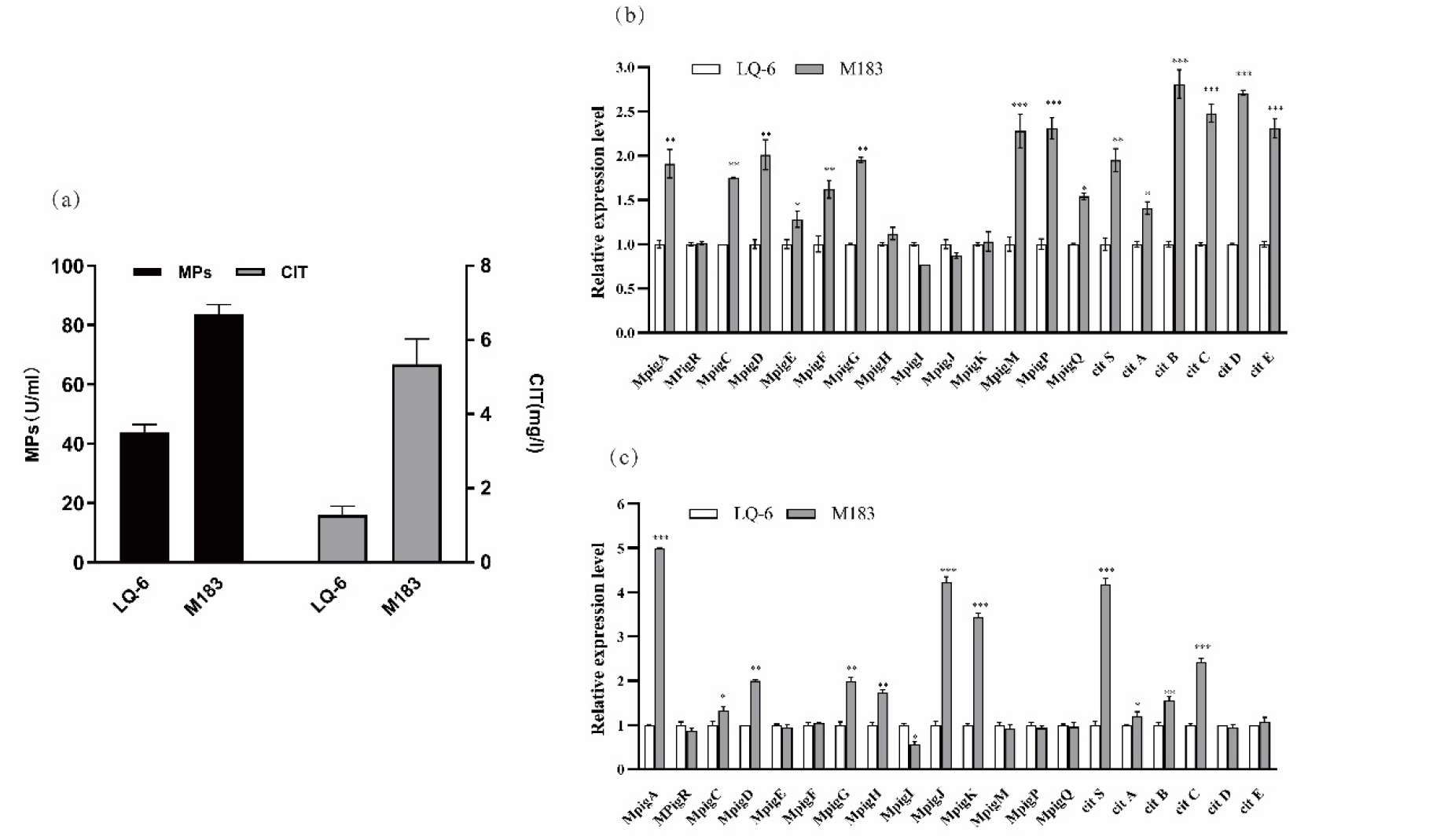
(a) Production of MPs and citrinin in SF of *M. purpureus* LQ-6 and *M. purpureus* CSU-M183 for 7 days. (b) Expression of genes related to MPs and citrinin biosynthesis of *M. purpureus* LQ-6 (control) and *M. purpureus* CSU-M183 at 4th day. (c) Expression of genes related to MPs and citrinin biosynthesis of *M. purpureus* LQ-6 (control) and *M. purpureus* CSU-M183 at 7th day

**Fig. 5.**
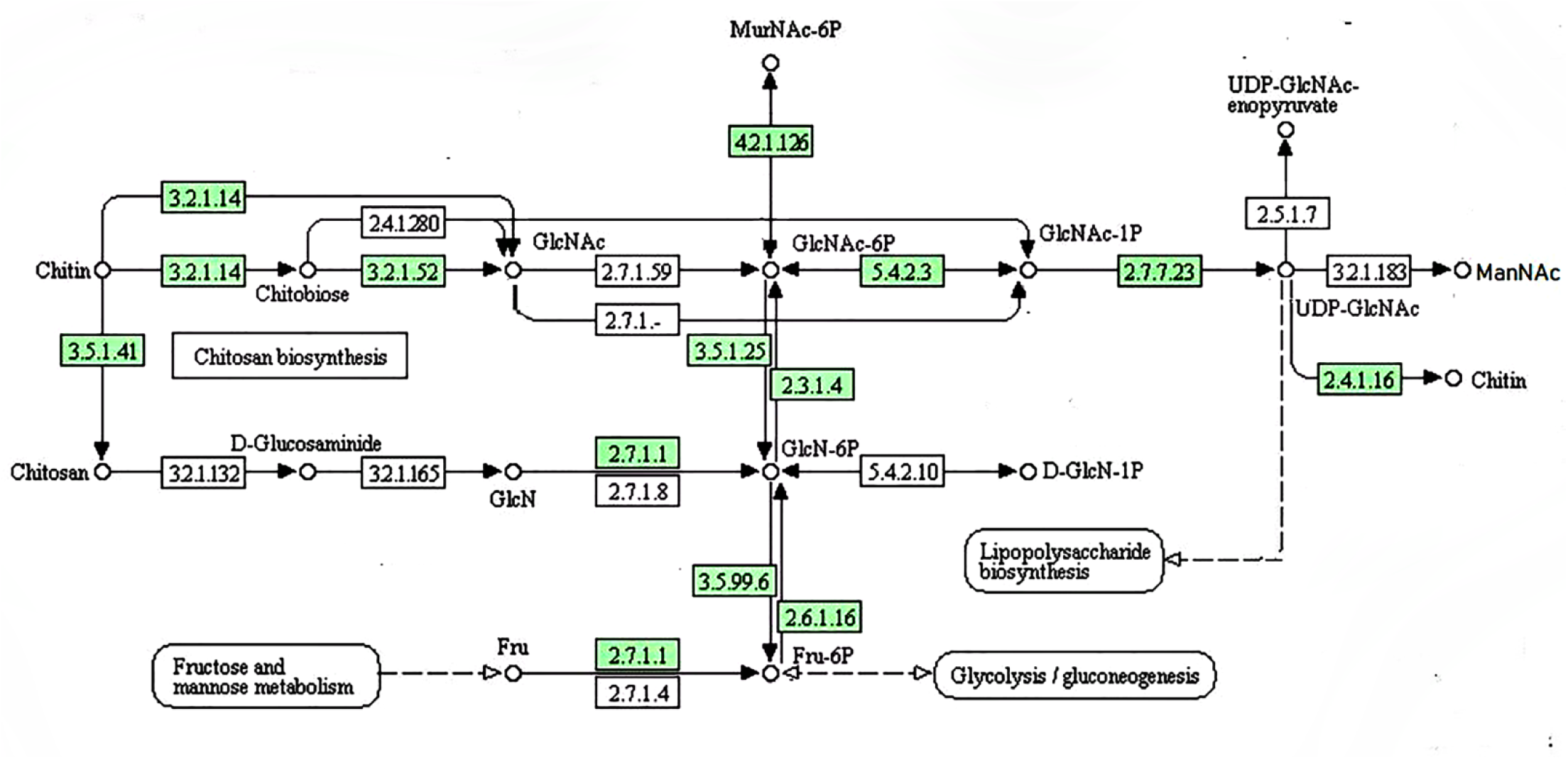
Prediction of chitin biosynthetic pathway in *M. purpureus* CSU-M183 genome, and the corresponding enzymes involved in each bioconversion step are shown in green

### 3.4 Analysis of the chitin biosynthesis pathway

As the main component of the fungal cell wall, chitin is important for the morphology of fungi. Based on the homology of amino acid sequence, chitin synthetases can be divided into three categories (class I-III) in *Saccharomyces cerevisiae*, four (class I-IV) in *Candida albicans*, and seven (class I-VII) in filamentous fungi. The numbers of gene encoding chitin synthetase in various filamentous fungi are different, generally containing 6-10 genes encoding chitin synthetase.[55]

To date, information about chitin biosynthesis in *M*. *purpureus* has not been reported. To lay a foundation for the further study of morphological metabolism of *M. purpureus*, we analyzed the chitin biosynthesis of strain CSU-M183 and annotated the function of the relevant genes in the pathway. By matching the predicted chitin biosynthesis-related enzymes in CSU-M183 strain genome with the KEGG database, the biosynthetic pathway of chitin in *M*. *purpureus* was identified. As shown in Fig. 4, phosphoacetylglucosamine mutase (PGM3) [EC:5.4.2.3] (encoded by gene g4907) converts N-acetyl-D-glucosamine 6-phosphate (GlcNAc-6P) to N-acetyl-alpha-D-glucosamine 1-phosphate (GlcNAc-1P), which is then dephosphorylated by UDP-N-acetylglucosamine diphosphorylase (UAP1) [EC:2.7.7.23] (encoded by gene g6630) to yield UDP-N-acetyl-D-glucosamine (UDP-GlcNAc). Moreover, chitin synthase (*chs*1) [EC:2.4.1.16] (encoded by genes: g872, g920,g3078, and g5640) converts UDP-GlcNAc to chitin. Additionally, the other genes encoding the important enzymes in the biosynthetic pathway of chitin were annotated, such as N-acetylmuramic acid 6-phosphate etherase (murQ) [EC:4.2.1.126] encoded by gene g1905, chitinase [EC:3.2.1.14] encoded by genes g3222, g6372 and g1142, and glucosamine-phosphate N-acetyltransferase (GNPNAT1, GNA1) [EC:2.3.1.4] encoded by gene g2832.

Class III chitin synthases only exist in the cell wall of fungi with high chitin content and are essential for regulating the mycelial aggregation morphology of fungi. Gene *chs*B in *Aspergillus fumigatus* plays an important role in cell wall biosynthesis, hyphal growth, and asexual reproduction.[56] To provide genetic resources for further studies, we mainly identified the gene *chs*B (gene g4739, 3243 bp) encoding class III chitin synthase in *M. purpureus* CSU-M183.

Generally, the specific characteristic of their SF is the aggregation of mycelia that are affected by environmental conditions, leading to different rheological properties of the fermentation broth. Such changes affect affect the transfer of mass, heat, and momentum, as well as the biosynthesis and production efficiency of target products. Moreover, the morphology of hyphae is closely related to the biosynthesis of secondary metabolites, and changes in the mycelium morphology of *Monascus* can regulate the level of secondary metabolites.[17, 24] As the main component of the fungal cell wall, chitin affects the mycelial morphological changes such as apical extension, branch growth, and differentiation. Blocking the biosynthetic pathway of chitin inevitably changes the mycelial aggregation and regulates metabolic pathways of target products. With the rise of the SF technology of *Monascus*, the effects of mycelial morphology on the biosynthesis of secondary metabolites in the fermentation process have attracted much attention. However, research on the metabolic engineering of *Monascus* morphology is still in the blank stage. In this article, we commented the strategies for morphological regulation of filamentous fungi, and discussed the impact of calcium signal transduction and chitin biosynthesis on apical hyphal growth and mycelial branching. Furthermore, based on the WGS analysis of strain *M*. *purpureus* CSU-M183, we will use genetic engineering technology to disturb the chitin biosynthesis of *M*. *purpureus* CSU-M183, change the mycelial aggregation morphology in the process of SF, regulate the biosynthesis of secondary metabolites, and clarify the molecular mechanism of the regulation of morphological on secondary metabolism using genetic engineering technology and histochemical correlation analysis.

## 4. Conclusion

The genomic information of *M. purpureus* CSU-M183 reported here can serve as a reference genome for *Monascus* genomics research. We predicted secondary metabolites BGCs and the chitin biosynthetic pathway in the genome of *M*. *purpureus* CSU-M183, and confirmed that ARTP induced significantly the relative expression of genes on MPs and citrinin biosynthetic gene clusters in *M. purpureus* CSU-M183 by RT-qPCR. Besides, we annotated and classified the chitin biosynthesis genes of *M. purpureus* CSU-M183, which offer a strategy of morphological metabolitic engineering. In conclusion, we provided genomic resources for further biological studies on the regulation of *Monascus* morphology on the biosynthesis of secondary metabolites.

## Author contributions

JL conceived and designed research. SZ and XZ annotated and analyzed the draft genome of *M. purpureus* CSU-M183. QL collected the samples. SZ and JL co-write the manuscript. All authors read and approved the final version of the manuscript.

## Acknowledgments

We would also like to thank Editage for English language editing. This work was supported by the National Natural Science Foundation of China (No. 32101906), Natural Science Foundation of Hunan Province (No. 2021JJ31146); Open Project Program of the Hunan Provincial Key Laboratory of Food Safety Monitoring and Early Waring (No. 2021KFJJ02), Education Department of Scientific Research Project of Hunan Province (No. 20B619) and Education Department of Postgraduate Research and Innovation Project of Hunan Province (No. CX20210863, CX20210898)

## Data availability

The complete genome sequence of *M. purpureus* CSU-M183 has been deposited into the NCBI Genbank database with an accession number of JAACNI000000000. The BioProject and BioSample information are available at PRJNA599556 and SAMN13759458, respectively.

## Compliance with ethical standards

## Ethical approval

This article does not contain any studies with human participants or animals performed by any of the authors.

